# Spatial Capture-Recapture with Partial Identity: An Application to Camera Traps

**DOI:** 10.1101/056804

**Authors:** Ben C. Augustine, J. Andrew Royle, Marcella J. Kelly, Christopher B. Satter, Robert S. Alonso, Erin E. Boydston, Kevin R. Crooks

## Abstract

Camera trapping surveys frequently capture individuals whose identity is only known from a single flank. The most widely used methods for incorporating these partial identity individuals into density analyses discard some of the partial identity capture histories, reducing precision, and while not previously recognized, introducing bias. Here, we present the spatial partial identity model (SPIM), which uses the spatial location where partial identity samples are captured to probabilistically resolve their complete identities, allowing all partial identity samples to be used in the analysis. We show that the SPIM out-performs other analytical alternatives. We then apply the SPIM to an ocelot data set collected on a trapping array with double-camera stations and a bobcat data set collected on a trapping array with single-camera stations. The SPIM improves inference in both cases and in the ocelot example, individual sex determined from photographs is used to further resolve partial identities, one of which is resolved to near certainty. The SPIM opens the door for the investigation of trapping designs that deviate from the standard 2 camera design, the combination of other data types between which identities cannot be deterministically linked, and can be extended to the problem of partial genotypes.

## 1. Introduction

The inferential goal of capture-recapture studies is to estimate population density, *D*, or abundance, *N*, in the presence of imperfect detection. Individuals are either naturally or manually marked and subjected to repeated capture attempts in order to estimate their capture probability and thus *D* or *N*. Generally, capture-recapture models for wildlife species regard the individual identity of each capture event as known; however in practice, the identities of individuals for some capture events can be ambiguous or erroneous. In live-capture studies, tags can be lost. In camera trapping studies, researchers often obtain partial identity samples – left-only and right-only photographs that cannot be deterministically linked. In genetic capture-recapture studies, partial genotypes and allelic dropout can lead to partial identification or misidentification, respectively. Statistical models have been developed to address the problem of imperfect identification in live capture using double tagging (e.g. Wimmer et al., 2013) and in camera trap and genetic capture-recapture studies by regarding the complete identification of partial or potentially erroneous samples as latent and specifying models for both the capture-recapture process and the imperfect observation process conditional on the capture (e.g. McClintock et al., 2013; Bonner and Holmberg, 2013; Wright et al., 2009). However, relatively little attention has been paid to one of the most important determinants of sample identity–the spatial location where it was collected. The identity of ambiguous samples should more likely match the identity of other samples collected closer together in space than those collected further apart and this information can be used to model the observation process and aid in the determination of sample identity.

The information about identity contained in the spatial location of samples has been used in two recent spatially-explicit capture-recapture (SCR) models where there is no other information about identity for some or all samples. Chandler and Clark (2014) probabilistically associate unidentified detections or counts to individuals identified by mark-recapture using their spatial location and a latent SCR model and Chandler et al. (2013) consider the situation where 100% of the samples are of unknown identity and use the spatial location of samples in combination with a latent SCR model as the basis for estimating density from such data (see Fewster et al., 2016, for an alternative model for spatially correlated unidentified counts). Further, spatial mark-resight models (e.g. Sollmann et al., 2013a) use the spatial locations of capture to resolve the uncertain identities of unmarked and sometimes marked individuals. In this case, mark status constitutes a partial identity. Here, we address the use of sample location to probabilistically resolve partial identities in camera trapping studies (i.e. single flank photographs) where complete identities are derived from two flanks.

Camera traps (remotely triggered infra-red cameras) have become an established method for collecting capture-recapture data for a wide range of species, especially those that are individually-identifiable from natural marks found on both flanks of the animal (termed “bilateral identification” by McClintock et al., 2013). Camera trapping studies typically allow capture-recapture data to be collected over longer periods of time and across larger areas than is feasible using live capture, leading to more captures of more individuals and thus more precision for population parameter estimates such as density (Kelly et al., 2012). These characteristics are especially advantageous when studying animals existing at low densities, such as large carnivores. However, even when using camera traps, researchers have found it difficult to achieve adequate precision for parameter estimates of low density populations, so any innovations in statistical methodology that can improve statistical efficiency, such as allowing unidentified or partial identity samples to be included in the analysis, are of broad practical interest.

Because animal markings are usually bilaterally asymmetric, researchers need to simultaneously photograph both flanks of an individual at least once during a capture-recapture study in order to obtain a complete identity (McClintock et al., 2013) and this is the reason the majority of camera trap studies deploy two cameras at each trap station. Given a single simultaneous both-side capture, all of the capture events for an individual can be combined into an capture history–the record of whether or not it was captured on each trap and occasion. For individuals that are never photographed on both flanks simultaneously, left-only and right-only photographs cannot be deterministically assigned to a single individual. These partial identity individuals can be linked across occasions using either their left-only or right-only captures, but it is not known which, if any, of these left-only and right-only partial identity capture histories are the same individuals. Single-sided photographs can occur in the standard double camera trap design if one camera is not triggered or has malfunctioned, one photograph is blurry, or the animal is photographed at an angle or position that only permits identification of a single flank. While less common in capture-recapture studies, the use of single camera trap stations can usually only produce single-sided photographs, none of which can be deterministically linked without supplemental information, such as dual-flank photographs from a live capture event (e.g. Alonso et al., 2015).

Including both left-only and right-only capture events in a single capture history may often result in instances where the capture events for a single individual are erroneously split across two individuals, one coming from the left-only captures and the other from the right-only captures. Therefore, researchers have typically discarded some of the single-sided captures from analysis (McClintock et al., 2013). If only single camera trap stations are used, left-only and right-only capture histories can be constructed. If at least some double camera trap stations are used, the left- and right-only captures can be linked to the complete identity individuals that were captured on both sides simultaneously at least once during the survey. In these scenarios, two capture histories are usually constructed – all capture events for the complete identity individuals are supplemented by either the left-only capture events or right-only capture events of the partial identity individuals. The most common approach is to analyze a single side data set and the chosen side is usually the one with more captured individuals or capture events (e.g. Kalle et al., 2011; Nair et al., 2012; Srivathsa et al., 2015; Wang and Macdonald, 2009). This process introduces two forms of bias that to our knowledge have not been identified in the literature. First, if the data set with more captured individuals is always the one selected for analysis, positive bias is introduced because the likelihood does not condition on this selection process. Second, linking all three capture types for the complete identity individuals introduces individual heterogeneity in capture probability and thus negative bias because the observed captures disproportionately come from the individuals with the highest capture probabilities (the complete identity individuals), leading to an overestimate of capture probability which in turn leads to an underestimate of abundance (Otis et al., 1978). To see how this process introduces individual heterogeneity, if *p_B_*, *p_L_* and *p_R_* are the probabilities of being captured on both sides, left-side only, and right-side only respectively, complete identity individuals will have a capture probability of *P*(*B*∪*L*∪*R*) = 1 − (1 − *p_B_*) (1 − *p_L_*)(1 − *p_R_*) *> p_L_* or *p_R_* by the monotonicity property of probability. A second approach that avoids the introduction of individual heterogeneity is to ignore the fact that the left and right side photos from a simultaneous capture belong to the same individual, average the density estimates from both single side analyses, and derive a joint standard error assuming independence. This method is proposed by Wilson, Hammond and Thompson (1999); however, Bonner and Holmberg (2013) point out that assuming independence between the dependent data sets will lead to the underestimation of standard errors and below nominal confidence interval coverage. Methods that appropriately model the dependence between the data sets by accounting for the imperfect identification process are thus required to produce unbiased estimates with appropriate measures of uncertainty.

Two recent papers (McClintock et al., 2013; Bonner and Holmberg, 2013) have extended the Latent Multinomial Model (LMM) of Link et al. (2010), originally applied to genetic capture-recapture with misidentification, to allow the complete and partial identity samples to be modeled together while accounting for the uncertainty in identity of the partial identity samples. Both papers show that the uncertainty stemming from the imperfect observation process is more than offset by the gain in precision from using all the capture events, leading to a net increase in precision of abundance (McClintock et al., 2013) and survival (Bonner and Holmberg, 2013) estimates, at least for the scenarios considered for simulation. While the MCMC-based LMM accounts for the uncertainty in identity by sampling from latent true capture histories that are consistent with the observed capture histories, this approach does not use the information about where samples were collected. However, using the spatial information associated with samples can reduce the frequency with which we propose and accept true capture histories that combine samples with locations that are unlikely to be from the same individual based on the movement characteristics of the species under consideration. For example, consider 3 samples (A, B, C) of unknown identity of a mesocarnivore with a typical home-range area of 4 km^2^. If samples A and B are 6 km distant, but samples A and C are only 1 km distant, then it is more likely that samples A and C are the same individual than samples A and B. SCR models are a natural framework for dealing with uncertain identity in capture-recapture models because they involve an explicit description of how the spatial organization of individuals interacts with the spatial organization of traps or other sampling devices. Therefore, we propose a spatial partial identity model (SPIM) that uses the spatial information associated with each photograph in camera trap studies to jointly model simultaneous, left-only, and right-only photographs while accounting for the uncertain identity of partial identity samples within the SCR framework. We apply this model to two data sets – one from a double camera station study of ocelots in Belize and one from a single camera station study of bobcats in southern California.

## 2. Methods – Model Description

We assume that individual activity centers ***s****_i_* are distributed uniformly across a continuous, two-dimensional state space 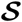 according to 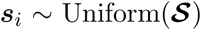 (but see Borchers and Efford, 2008; Reich and Gardner, 2014; Royle, Fuller and Sutherland, 2016, for alternative specifications). This state space is a rectangular or polygonal user-defined region inhabited by the population. Next, let ***x*** be a *J* × 3 matrix for the *J* traps, with the first two columns containing the *X* and *Y* coordinates of the traps and the third column containing the number of cameras deployed at each station. We define events *m* ∈ (*B, L, R*) to correspond to both-side simultaneous capture, left-only capture, and right-only capture, respectively. We assume a partially latent binomial capture process such that for the *m^th^* capture type, 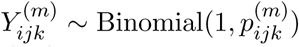 with 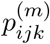 being the capture proobability of individual *i* at trap *j* on occasion *k* (e.g., day of a camera trapping study) for event type *m*. The true, partially latent capture history is then the set of binomial frequencies 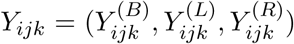 and the observed capture history is the set of binomial frequencies 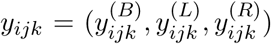. The dimensions of the three binomial frequencies are *n_B_* × *J* × *K*, *n_L_* × *J* × *K*, and *n_R_* × *J* × *K*, respectively, with, *K* being the total number of trap nights and *n_m_* being the number of individuals for which at least one *m* event was observed. Both true and observed capture histories are augmented up to dimension *M* × *J* by adding *M* − *n_B_*, *M* − *n_L_* and *M* − *n_R_* rows of all zero capture histories (see Royle, Dorazio and Link, 2007; Royle, 2009, for a complete description of data augmentation in capture-recapture models). A vector of *M* partially latent indicator variables **z** is introduced to indicate which individuals are in the population with *z_i_* ~ Bernoulli(*ψ*), inducing the relationship *N* ~ Binomial(*M, ψ*). Therefore, population abundance, *N* is a derived quantity obtained at each MCMC iteration by 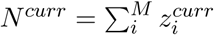 and so is population density, 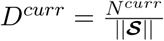.

Conditional on the partially latent ***Y***, the detection process is that of a typical SCR model, except that the capture probabilities depend on the capture type and number of cameras at a trap. We assume a Gaussian hazard detection function. Because double cameras are typically positioned to fire together in order to get a both-side capture, it is unlikely that each camera is independent. Rather than model the capture probabilities of each of the two cameras with some correlation between cameras, we specify different capture probabilities for both-side captures and single-side captures, assuming independence between the both, left, and right-side capture processes. We write the detection function for capture type *m* as
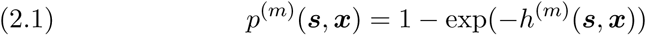

with *p*^(*m*)^(***s***, ***x***) being the probability of capture for an individual with activity center ***s*** at trap location ***x*** and
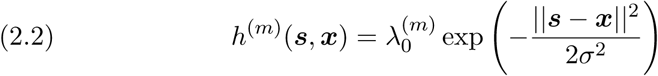

with 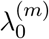 being the expected number of detections for capture type *m* for an activity center located at the same location as a trap and *σ* being the spatial scale parameter that determines how quickly capture probability declines with distance between an activity center and a trap. Because we do not expect any systematic difference in the probability of detecting one flank over the other, we will set 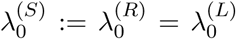 where *S* indicates a single side capture. For single camera stations, we have the single-side detection function *p*^(1*S*)^(***s***, ***x***) for both L and R captures. At double camera stations, the single side capture probability is
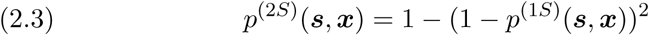

because there are now two ways to photograph a single side (camera 1 or 2). Note that *p*^(.)^(***s***, ***x***) here corresponds to equation 2.1 for a single side capture. *B* captures can only occur at double camera stations and so we introduce 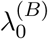 as the expected number of both-side observations for an activity center located at the same location as a trap. Finally, the single and both-side capture probabilities for each individual at each trap depends on the number of cameras deployed following 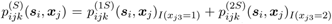 and 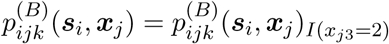.

If we knew the complete identities of all captured individuals, we could construct the true capture history ***Y*** from the observed capture history ***y*** by reordering the rows of ***y***^(*L*)^ and ***y***^(*R*)^. Thus, the key idea behind the SPIM is that we can sample the latent true capture histories by simply reordering the *i* indices of ***y***^(*L*)^ and ***y***^(*R*)^ accordingly. To do this, we define ***Y***^(*B*)^ and ***y***^(*B*)^ to be in the correct order of identity, corresponding to the order of **s** and **z**, and introduce identity vectors to aid in updating the latent *i* indices of the true capture history. We specify the known identity vector 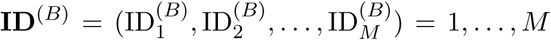. Then, we introduce partially-latent identity vectors 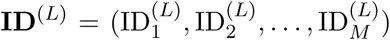 and 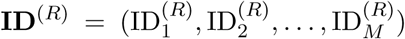 indicating which 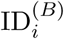 each 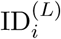 and 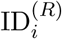 correspond to. For example, if the values of 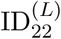 and 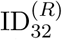 are 28, the left and right capture histories for the 28th individual in ***Y***^(*B*)^ and ***y***^(*B*)^ are stored in the 22^nd^ and 32^nd^ *i* indices of ***y***^(*L*)^ and ***y***^(*R*)^, respectively. On each MCMC iteration, we construct latent true capture histories ***Y*** from the latent identities in **ID**^(*L*)^ and **ID**^(*R*)^ after swapping some of the identities and associated partial identity samples between activity centers. This process produces posterior distributions for the SCR parameters that account for the uncertainty in identification of the partial identity samples.

To prevent samples from the complete identity individuals from being swapped, we define **c** to be an *n* × 1 indicator vector with entries 1 if the complete identity of individual *i* is known, whether from a *B* event at some point during the study or from auxiliary data, and 0 otherwise. In all cases of c*_i_* = 1, individual identities are complete and ***Y***^(*m*)^ = ***y***^(*m*)^ for *m* ∈ (*B, L, R*). Conversely, the *i* indices of ***y***^(*L*)^ and ***y***^(*R*)^ with c*_i_* = 0 are partial identity samples and ***Y***^(*L*)^ and ***Y***^(*R*)^ are latent. For convenience, we jointly sort ***Y*** and ***y*** such that the 1,…, *n*_Complete_ c*_i_* = 1 individuals are the first individuals to occur in the true and observed capture histories, 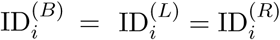 for the first *i* = 1,…, *n*_Complete_ individuals, and we only need to resolve the latent identities of the *i* = *n*_Complete+1,…, *M*_ *i* indices of the ***y***^(*L*)^ and ***y***^(*R*)^ observed data sets.

### 2.1. Methods – Trap Operation File

It is common for cameras to malfunction in camera trap studies. In typical SCR, this can be accommodated by modifying the capture process using ***L*** (dimension 1 x J), the row vector containing the number of trap nights each trap was operational, if working with the 2-D data matrix (individual × trap summed over occasions). If working with the 3-D data array (individual × trap × occasion), ***L***, the complete trap operation history matrix is then a matrix whose *jk^th^* element is 1 if trap *j* was operational on occasion *k* and 0 otherwise. Then, *y_ij_* ~ Binomial(*L_j_, p_ij_*), or *y_ijk_* ~ Binomial(1*, p_ijk_* × *L_jk_*). Traditionally, the trap operation file does not distinguish between having one or two cameras operational, despite the capture probability likely being higher when both traps are functional. In the SPIM, both side captures can only occur when two cameras are operational and the probability of a single side capture depends on whether one or two cameras are operational; therefore, the 3-D data array needs to be used to properly account for camera operation if there are two camera stations. Specifically, the single and both-side trap by occasion level detection probabilities for each individual are then 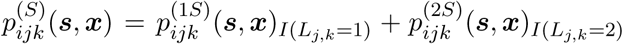 and 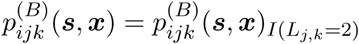, respectively.

### 2.2. MCMC Algorithm

We will describe the novel aspects of the MCMC algorithm here – see Appendix A for the complete description. The following are our uninformative prior distributions.

1. 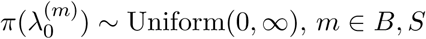
2. *π*(*σ*) ~ Uniform(0, ∞).
3. *π*(*ψ*) ~ Uniform(0, 1)
4. 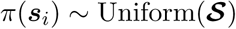

The joint posterior is then:

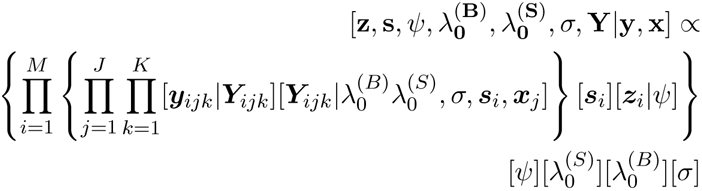

and we sample from this distribution using MCMC. The full conditional for ***Y****_i_*^(*m*)^ is:

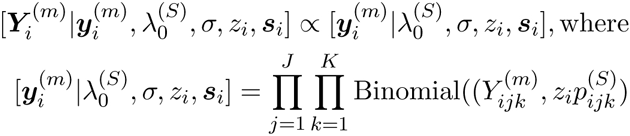

for *m* ∈ (*L, R*). While not part of the joint posterior, the ID vectors can be used to update ***Y*** and conditional on **ID**^(*L*)^ and **ID**^(*R*)^, we can construct a latent true capture history ***Y*** so our MCMC algorithm follows the standard SCR MCMC algorithm as described by Royle et al. (2013) with the additional step of updating **ID**^(*L*)^ and **ID**^(*R*)^ to produce a new latent true capture history ***Y*** on each MCMC iteration. On each MCMC iteration, we update both **ID**^(*L*)^ and **ID**^(*R*)^ by swapping *n_swap_* values of **ID**^(*B*)^ stored in **ID**^(*L*)^ and **ID**^(*R*)^. We first update **ID**^(*L*)^. We need to identify the correctly ordered indices **ID**^(*B*)^ at which to swap the value of **ID**^(*L*)^, mapping **ID**^(*L*)^ to **ID**^(*B*)^. We then identify the candidate set of **ID**^(*B*)^ individuals that do not correspond to complete identities (c*_i_* = 0) and who are currently in the population (*z_i_* = 1). Next, we choose a focal candidate *v* to swap the value of 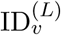 with equal probability across the candidate set. Because proposals that combine candidates whose activity centers are far apart will almost always be rejected, we apply a distance-based criterion to rule out improbable combinations, thus raising acceptance rates. To do this, we calculate the Euclidean distance between the current activity center of the focal candidate *v* and the activity centers of all other individuals in the candidate set. We then identify the set of possible candidate individuals to exchange values of 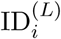 with the focal candidate by identifying which candidate individual activity centers are within a distance threshold, *d_max_*, of the focal individual’s activity center. From this reduced candidate set of size *n_forward_*, we randomly select individual *w* with equal probability 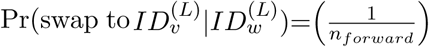 across the remaining candidates and the focal and selected candidate exchange values of 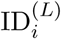. Because this proposal process is not symmetric, we repeat it in reverse to obtain *n_reverse_*, with the probability of choosing this candidate being 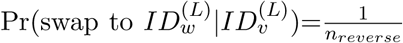. We recompute the proposed true capture history 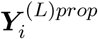 for *i* ∈ *v*, *w* and accept the proposal with probability
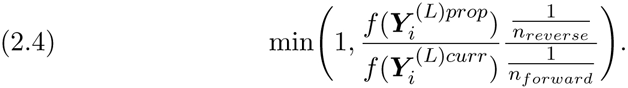

where 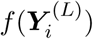 is the SCR observation model likelihood. This process is then repeated to update **ID**^(*R*)^ and thus, ***Y***^(*R*)^.

### 2.3. Methods –”Pragmatic” Estimators

We will consider the most common estimator used in practice based on choosing the single side data set with the most captured individuals and combining it with the both side data set, if available, and analyzing the resulting data set with a traditional null SCR model (fixed *λ*_0_ and *σ*). Because choosing the best single side introduces a positive bias, the second estimator will choose a random side to be combined with the both side data set, if available. We will refer to these estimators as the “best-side” and “random-side” estimators on single camera trapping arrays and “both-plus-best-side” and “both-plus-random-side” estimators on double camera and hybrid trapping arrays.

## 3. Application 1: Dual camera station trapping array targeting ocelots

This data set comes from a long-term, multi-site felid study in Belize conducted from 2008 to the present for which an analysis has not yet been published. The study targeted jaguars, pumas, and ocelots, but due to their smaller size and more nocturnal activity patterns, the probability of simultaneously photographing ocelots on both flanks was relatively low, leading to several ambiguous single-sided capture histories within any given year. Because this is a multi-year study, the complete identities of some individuals within any given year are known from other years, but we will use a single data set in isolation to model the more typical single year survey. This specific data set was collected in the Rio Bravo Conservation Management Area, Belize, in 2014. The trapping array (Figure 1a) consisted of 26 dual camera stations with a mean spacing of 1.96 km and the survey lasted 98 days (July 20 - October 25), resulting in 1796 trap nights with 2 cameras operational and 425 trap nights with a single trap operational due to malfunction. Sex could be determined from the photographs for all individuals except for one individual that was captured a single time. Eight individuals (5 male, 3 female) were captured on both flanks simultaneously at least once during the experiment producing complete identities and another (male) was captured on both flanks at a single camera station in short succession such that it was improbable that both sides did not belong to the same individual. This individual’s identity was considered complete and the capture was recorded as a left-side capture, chosen randomly. This was done because a single camera was operational during this event and our model does not allow a both-side capture to occur when a single camera is operational and recording the event as both a left and right capture would violate the independence assumption between the capture processes. There were nine partial identity left-side capture histories (1 male, 7 female, and 1 unknown) and 12 partial identity right-side capture histories (5 male and 7 female). From other years, it is known that 5 of the partial identity left capture histories belong to individuals recorded in the right capture histories. Overall, there were 10 both-side captures, 30 left-side captures, and 48 right-side captures. The spatial distribution of captures for partial identity individuals can be seen in Figure 1a.

**Fig 1:**
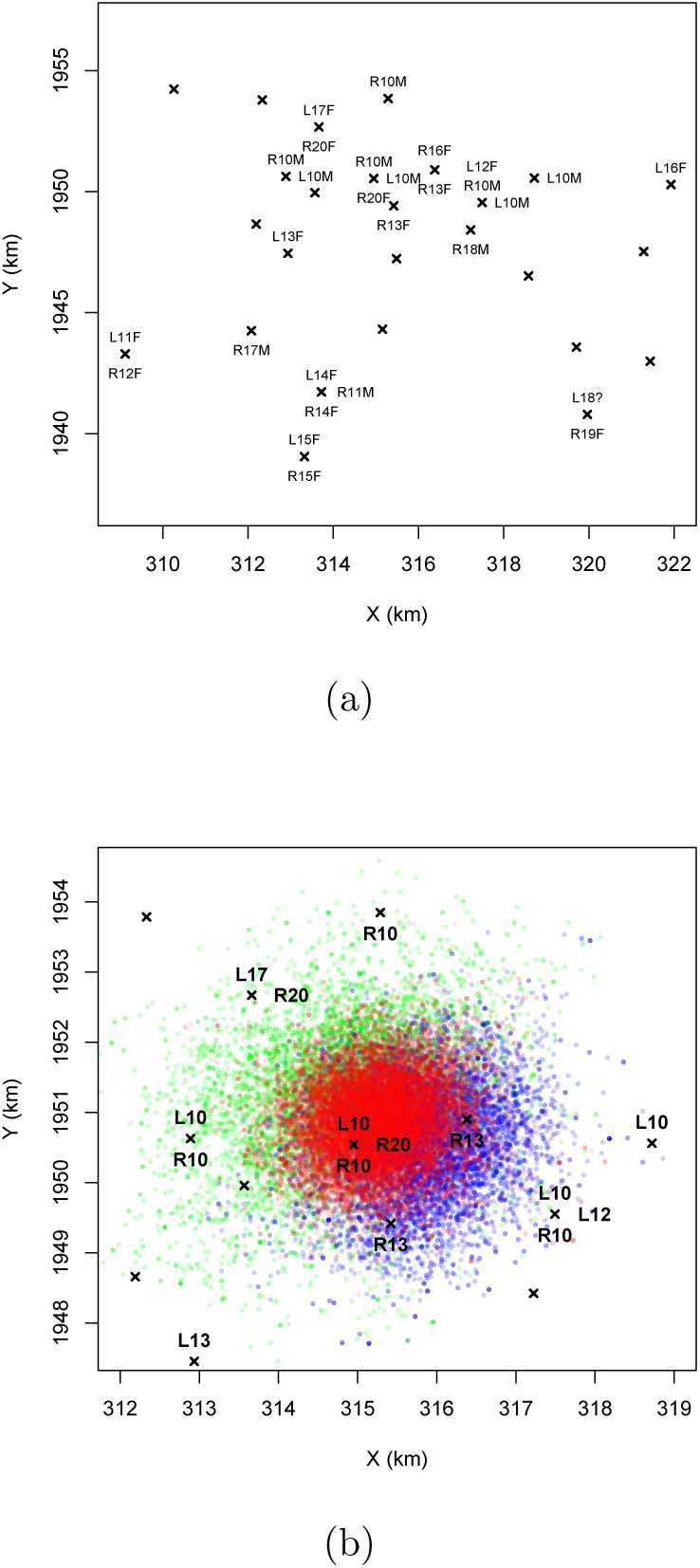
(a) Capture locations for partial identity samples in the ocelot data set. R and L indicate right and left, respectively and M, F and ? indicate male, female, and unknown, respectively. (b) The posterior distribution for L10 and R10 when they are correctly matched (red), for L10 when not matched to R10 (green) and for R10 when not matched to L10 (blue). When L10 is not matched to R10, it mostly matches with R13 and R20. When R10 is not matched to L10, it mostly matches L12, L13, and L17. These results are from the model not using sex information.

We analyzed the complete data set, the male-only data set, and the female-only data set. Knowing the sex of almost all individuals provides us the opportunity to exclude matching partial identity samples of different sexes; however, this information is not observable from camera trap photographs for many species. To model this more common situation, we first analyzed the full data set without using the sex covariate. Then, we used the sex covariate to exclude matches between sexes to model either the situation where sex is known from photographs or that of a species living at a lower density than this population of ocelots. Our model could be modified to allow matches based on categorical covariates such as sex while sharing the same density and detection function parameters; however for convenience, and because male and female ocelots likely do not share the same *σ* or *D* (M. Kelly, unpublished data), we analyzed the male and female-only data sets separately. This is conceptually equivalent to formally including a sex covariate in the SPIM and allowing all parameters to vary by sex. For all three data sets, we fit the SPIM to the full data set and traditional SCR models to data sets that augmented all captures for the complete identity individuals (both, left, and right) by either the left or the right partial identity capture histories. For all models, we ran one chain for 35K iterations, discarding the first 5K, and in the SPIMs, we set *d_max_* to 3 km and *n_swap_* to 10. Based on the simulations of double camera trap station surveys, we expected the SPIM estimates to be slightly less precise, but slightly larger due to the individual heterogeneity introduced by the traditional manner of combining the three data sets.

The results in Table 1 largely matched our expectations. The density estimates of the SPIM were higher than the mean of the two SCR0 estimates by 21, 32, and 31% for the total, male, and female data sets, respectively. The right side data set was the “best-side” data set and it produced an estimate closer to the SPIM, which matches the simulation results. 95% HPD intervals were slightly narrower using traditional SCR in 4 of the 6 possible comparisons and slightly narrower using the SPIM in the remaining 2. *σ* estimates for males were higher than for females and did not vary widely among the three methods of analysis. Adding the posteriors for *N* from the male and female only models produced an estimate of 40 (29 - 58), which was one unit narrower than the SPIM not including sex information, despite including three extra parameters and excluding the individual of unknown sex. Overall, the SPIM provides more optimistic density estimates, that according to the simulations, should be closer to the truth with credible intervals that can provide nominal frequentist coverage or offer more accurate Bayesian interpretations, and removes the need to interpret two sets of estimates. For the complete data set, the SPIM took 146 minutes to run on a laptop with a 2.7 GHz Intel I7 processor.

**Table 1.** Parameter estimates for the ocelot data set using either the Spatial partial identity model (SPIM) or the standard spatial capture-recapture model (SCR0) on either the both plus right side data set or both plus left data set. Density is in units of individuals per 100 km^2^.

| Sex | Model | $p_0^S$ | $p_0^{(B)}$ | $p_0$ | $\sigma$ | N (95% CI) | D (95% CI) | CI width |
| --- | --- | --- | --- | --- | --- | --- | --- | --- |
| Both | SPIM | 0.005 | 0.003 |  | 2.00 | 42 (29 – 59) | 7.33 (5.09 – 10.36) | 5.27 |
|  | SCR0-B+R |  |  | 0.015 | 2.05 | 39 (27 – 56) | 6.87 (4.74 – 9.83) | 5.09 |
|  | SCR0-B+L |  |  | 0.015 | 2.25 | 30 (20 – 44) | 5.27 (3.51 – 7.73) | 4.21 |
| Male | SPIM | 0.005 | 0.004 |  | 2.40 | 14 (11 – 23) | 2.63 (1.93 – 4.04) | 2.11 |
|  | SCR0-B+R |  |  | 0.015 | 2.45 | 15 (11 – 26) | 2.72 (1.93 – 4.57) | 2.63 |
|  | SCR0-B+L |  |  | 0.022 | 2.39 | 8 (7 – 16) | 1.41 (1.23 – 2.81) | 1.58 |
| Female | SPIM | 0.005 | 0.002 |  | 1.21 | 24 (14 – 40) | 6.18 (3.56 – 10.18) | 6.61 |
|  | SCR0-B+R |  |  | 0.016 | 1.24 | 19 (12 – 35) | 4.89 (3.05 – 8.90) | 5.85 |
|  | SCR0-B+L |  |  | 0.007 | 1.71 | 18 (10 – 38) | 4.57 (2.54 – 9.67) | 7.12 |

The posterior distributions of sample identity for the partial identity samples provide interesting anecdotes about how both spatial location and a categorical covariate can individually, and in combination, inform sample identity. In the model not using information about individual sex, the 5 partial identity individuals in the left and right data sets that were known to be the same individuals from other surveys were assigned higher posterior probabilities of being the same individual than any other partial identity individuals (data not shown). Using location alone, these probabilities ranged from 0.23 - 0.74 and when adding the information about sex, they increased to 0.59 - 0.99 (Table 2). The tenth left and right partial identity histories, L10 and R10, had a high probability of (correctly) being the same individual with or without using the sex information (0.74 and 0.99, respectively). In Figure 1b, it can be seen that L10 was captured in 4 locations and R10 in 3 locations with roughly the same mean capture location. Incorrectly matching R10 with L12 pulls the combined mean capture location to the east, and incorrectly matching R10 with L13 pulls it to the south. Incorrectly matching L10 with R13 pulls the combined mean capture location to south and slightly to the east and matching L10 with R20 pulls it to the east and slightly to the north. These observations are reflected in the posterior distribution for the activity center of these two partial identity samples decomposed into the MCMC iterations when they were correctly matched and when they were not. When including sex information, we know that R10 (male) cannot match either L12 or L13 (females) and L10 cannot match R13 or R20 (females). This only leaves augmented individuals for L10 and R10 to incorrectly match and two augmented individuals, uncaptured by definition, with activity centers in the middle of the trapping array are very improbable. Therefore, the model assigns a 0.99 probability that L10 and R10 are the same individual when sex is considered. Conversely, L11 and R12 with no nearby same sex matches have a lower posterior probability of being the same individual (0.60) because they can plausibly be assigned to augmented individuals living off of the trapping array that were never right or left-captured, respectively.

**Table 2.** Posterior probabilities that left and right ocelot samples are from the same individual for individuals that were determined to be the same from data collected in other years.

| L ID | R ID | Pr(L ID = R ID) |  |
| --- | --- | --- | --- |
|  |  | Sex Unknown | Sex Known |
| 10 | 10 | 0.74 | 0.99 |
| 11 | 12 | 0.28 | 0.59 |
| 12 | 13 | 0.42 | 0.72 |
| 14 | 14 | 0.23 | 0.63 |
| 15 | 15 | 0.38 | 0.70 |

## 4. Application 2: Single camera station trapping array targeting bobcats

This data set comes from a study of bobcats in southern California that has been analyzed using both non-spatial partial identity models (PIM, McClintock et al., 2013; McClintock, 2015) and hybrid mark-resight models (Alonso et al., 2015) that combine mark-resight and capture-recapture for the unmarked, but individually-identifiable individuals. The trapping array consisted of 30 single camera stations with a mean spacing of 1.63 km operated over 187 days, producing 4669 trap nights and 109 left-only or right-only capture events of 23 left-side and 23 right-side individuals. Twenty-seven bobcats were GPS-collared, marked, and photographed on both sides at capture so their left- and right-side capture histories could be linked and 15 of these individuals were later photographed at camera traps. See Alonso et al. (2015) for a full description of the survey.

Following McClintock et al. (2013) and Alonso et al. (2015), we analyzed the data set in two ways. First, we analyzed the data set using the 15 complete identities obtained from the live captures to compare performance to the PIMs in McClintock (2015) and the hybrid mark-resight estimators in Alonso et al. (2015). While the hybrid mark-resight estimator makes use of the number of marked individuals in the population that were not recaptured, we did not constrain our MCMC sampler with this information so that a better comparison could be made to the PIM analyses that did not use this information and because the posterior density of N for the SPIMs placed negligible weight below the known number of individuals in the population during the survey (41). For the second analysis, we discarded the complete identities to model a single camera capture-recapture survey that did not have a live capture component. Because Alonso et al. (2015) found strong support for individual heterogeneity in the mark-resight models and both Alonso et al. (2015) and McClintock (2015) found moderate support for individual heterogeneity in capture-recapture models, we compare the SPIM to the PIM and mark-resight models with individual heterogeneity in capture probability. For each SPIM and SCR analysis, we ran one chain for 35K iterations, discarding the first 5K. For the SPIM models, we set *n_swap_*=10 and *d_max_* to 2 km. The SPIM models with and without the 15 complete identities took 57 and 53 minutes to run on a laptop with a 2.7 GHz Intel I7 processor, respectively.

Among the models using the 15 complete identity individuals, the most precise estimate was the hybrid mark-resight model using the right-side data set for the capture-recapture of unmarked individuals; however, the SPIM was more precise than the average of the left and right side analyses and removes the task of interpreting two estimates (Table 3). The conservative approach would be to interpret the least precise single side analysis, in which case the SPIM was 14% more precise than both the single-side hybrid mark-resight and SCR analyses. The SPIM was 66% more precise than the PIM with individual heterogeneity, which was considerably less precise than the classical M*_h_* single-side analyses discarding the 15 complete identities. When the 15 complete identities are discarded, the precision of the SPIM is only slightly reduced and is still 6% more precise than the least precise single-side hybrid mark-resight estimate and is 30% more precise than the least precise SCR estimate. The former suggests that there is a similar amount of information about density in the spatial location of captures on this single camera array as there is in knowing the marked status of 15 individuals and that the SPIM can remove the need for the live capture component of a study if the only goal is to mark individuals for mark-resight density estimation. While the SPIM appears to perform the most favorably on this data set compared to alternatives considered, we note that a definitive comparison would require a simulation study where the true parameter values are known and more than one survey can be conducted.

**Table 3.**
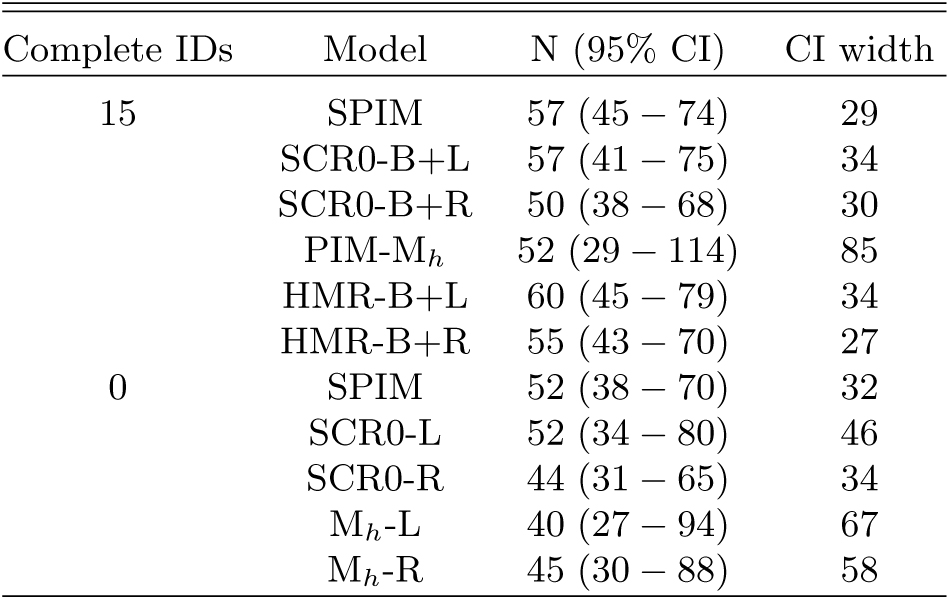
Population size estimates for the bobcat data set from the spatial partial identity model (SPIM), single side SCR analyses (SCR0), non-spatial partial identity models (PIM M_h_ McClintock, 2015), M_h_ hybrid mark-resight models (HMR Alonso et al., 2015), and M_h_ classical mark-recapture models (Alonso et al., 2015). The SPIM and single side SCR analyses are repeated both with (Complete IDs=15) and without (Complete IDs=0) the information from live-captured individuals.

## 5. Discussion

Our study has shown that the spatial locations where samples were collected provides information about individual identity and using this information in partial identity models can improve inference in camera trap studies. Further, the formal treatment of the number of cameras at trap stations allows for camera number and the spatial distribution of station types (1 or 2 cameras) to be considered when designing surveys. Simulations in Appendix B demonstrate that the SPIM estimator performs better than the best-side and random-side estimators, at least in the sparse data scenarios considered here. In general, the SPIM offers better performance gains in smaller populations, when there are fewer complete identity individuals, and when the percentage of individuals that have partial identities is higher. The performance gains in the hybrid designs was better than the all double designs because they produced fewer complete identities and a higher percentage of partial identities. In fact, the precision of the hybrid designs was not substantially lower than the all double designs, despite using only 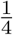 the number of double camera stations. This result suggest that hybrid designs could potentially be the best use of a fixed number of cameras – designs that to our knowledge are not currently being used. Another determinant of the ratio of partial to complete identity individuals is the ratio of 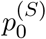 to 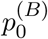. For example, the trapping array in the ocelot example also targeted jaguars which when photographed, are significantly more likely than ocelots to produce a complete identity because of their larger size, slower traveling speed, and less nocturnal activity patterns (M. Kelly unpublished data), perhaps reducing potential performance gains by using the SPIM.

The SPIM likely performs better on more regular, closely-spaced (relative to sigma) trapping arrays as investigated in the simulations. Partial identity samples on the interior of a regular, closely-spaced trapping array are more likely to be correctly matched than those on the edge of the trapping array or on a trapping array that is spaced more widely because it is less likely that an animal will only have a single side captured when it is surrounded by traps than if it is not. This can be seen in the ocelot example where the probability the right and left sample number 10 are the same individual is very high. In the model not including sex, each sample is never assigned to an augmented individual (an animal with the other side not captured) and when sex information is included, all other nearby partial identity samples are ruled out and the probability the samples match is estimated to be 0.99. This high certainty relies on the samples being on the interior of the trapping array in an area where the trapping array is roughly regular, because if these two samples do not match, there must be two augmented individuals living on the interior of the trapping array for each to match with and this is improbable. Conversely, left ID 11 is assigned to right ID 12 with probability 0.28 without sex information and 0.59 with sex information. This reduced certainty is mostly due to the partial identity samples being collected on the periphery of the array where augmented individual activity centers are much more likely to exist to be matched with. By the same argument, the SPIM should perform better on larger arrays where the ratio of interior to exterior array area is larger, given the same number of individuals are on the array. In our simulations, the best precision and MSE gains between the 6 x 6 and 8 x 8 arrays depended on the scenario, but we fixed *D* and so *N* varied by array size. Confirming this result requires further simulation.

As seen in the ocelot example, if an individual covariate aside from spatial location is available, the probabilities of correctly assigning the left ID to the correct right ID and vice versa can be considerably increased. We suspect this should in general increase precision for abundance and density by reducing the pool of potential matches for each partial identity sample. Indeed, in the ocelot example, when we added the male- and female-only posteriors for N, we slightly increased precision despite having modeled 3 additional parameters over the combined model and excluded the individual whose sex was not known. Reducing the set of potential matches should reduce the span of values of *p*_0_, *σ*, and number of captured individuals that are consistent with the data, increasing precision of abundance and density. We suspect the relative value of spatial location and other covariates depends on the degree they deterministically or probabilistically rule out potential matches. In general, knowing sex will rule out approximately half of the potential matches, while knowing spatial location on a large trapping array relative to *σ* should rule out a much higher percentage of matches. In the ocelot example, we took an ad hoc approach to using the sex information, but sex or other categorical covariates could formally be modelled either by ruling out inconsistent matches only between observed partial identity individuals, or by also modelling the category proportions (e.g. sex ratio) and updating the latent category values of the augmented individuals on each MCMC iteration.

A comparison of the SPIM to the non-spatial partial identity model of McClintock (2015) can be found in Supplement B. While the PIM estimator reliably decreased MSE, removed small sample bias, and increased precision in some scenarios, it reduced precision in the more data sparse scenarios we considered and offered only small precision gains in the presence of individual heterogeneity in capture probability. In general, we think individual heterogeneity in capture probability is difficult for the PIM to accommodate. Because the multinomial observation process (left, right, or both-side capture) is defined conditional upon capture, the likelihood that two partial identity capture histories are the same depends on how consistent their combined number of captures across capture types are with *p* and *N*. If all individuals can have their own *p*, the number of times the composite individual was captured becomes much less informative about identity. Because the left, right, and both-side capture processes in the SPIM are independent, the likelihood component for partial identity, single-sided capture histories does not depend on the combined number of capture events. Rather, the likelihood that two partial identity capture histories are the same depends on how consistent the combined *spatial distribution* of captures are with *p*_0_ and *σ*. Therefore, there should be less information about individual identity when there is individual heterogeneity in *σ*, and perhaps to a lesser extent, *p*_0_. Generalizations of the 2-flank SPIM to scenarios where partial identities cannot be categorized into types will require a model similar to the PIM estimator where the combined number of captures is informative about identity and hard to distinguish from individual heterogeneity. This problem arises in all SCR models with latent individual identities, such as spatial mark-resight and unmarked SCR, so the sensitivity of these models to individual heterogeneity in *p*_0_ and *σ* should be investigated.

One concern of using the SPIM over traditional SCR or the PIM is computational efficiency. We feel the computation demands of the SPIM are reasonable, at least for the low density scenarios where precision gains are the most needed. An R package to fit the SPIM is available at github.com/benaug/SPIM which includes code to fit the models in either R or Rcpp and RcppArmadillo (Eddelbuettel and François, 2011; Eddelbuettel and Sanderson, 2014), which is considerably faster. If a trap operation file is used and the 3-D data array must be used, the R analysis is much slower, but only slightly slower in Rcpp. In simulations with random trap failure (data not shown), ignoring trap failure reduced the estimates of 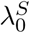 and 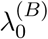, but not *N*, suggesting the use of the 3-D data array is not necessary, at least when trap failure is at random, but this warrants further investigation, To provide some benchmarks, we replicated scenario S9.6 on a laptop with a 2.7 GHz intel I7 processor, raising N to 100 with M=150. To run 35K MCMC iterations, it took 106.7 minutes in R and 10.3 minutes in Rcpp (~10x faster) with no trap file and the 2-D data matrix. Using the 2-D trap file and 3-D data array, it took 575.9 minutes in R and 12.6 minutes in Rcpp (~45x faster). Computation time can further be reduced using the semi-complete likelihood approach of King et al. (2015) which is currently being developed for the multimark package (McClintock pers. comm.) The longer reported run times for the bobcat and ocelot data sets are due to the use of polygonal, rather than rectangular state spaces, and reflect the computational demand of ensuring that activity center proposals falling outside of the continuous, many-sided state space are not accepted.

As previously recognized by Wright et al. (2009), another application where the spatial location of partial or potentially corrupted identity samples would be useful is in capture recapture studies using microsatellite markers. Wright et al. (2009) developed a non-spatial model that accommodated both partial genotypes and allelic dropout. In genetic capture-recapture studies, the spatial location where samples were collected is almost always recorded and could be used to resolve partial and potentially corrupted identities. The potential for improved inference is perhaps greatest for studies using genotypes from sources with low complete amplification rates due to small amounts of DNA or higher levels of degradation such as scat samples in tropical environments (e.g. Wultsch, Waits and Kelly, 2014); however, if these low quality samples are more likely to be erroneous, the misidentification process should be modelled. Unlike the camera trap observation model, the partial identity genetic samples have traditionally been completely discarded, suggesting that performance gains could be larger than seen here. One last potential DNA-based application is that researchers may choose to genotype fewer loci than necessary to determine a sample is unique in the population and model the resulting uncertainty in identity using the SPIM. This could either save project resources or allow more samples to be amplified for the same amount of resources. Since the information about identity in each loci comes with diminishing returns per additional loci, it is not clear that the better use of resources is to genotype fewer samples to a high level of certianty rather than to genotype many samples to a lower degree of information about identity. Finally, the SPIM could also be extended to combine any capture-recapture data types where identity cannot be resolved between methods. For example, Sollmann et al. (2013b) combined capture-recapture data from camera traps and scat samples by sharing *σ* between data sets. Using the SPIM, the latent structure (e.g. activity centers and **z**) could also be probabilistically shared. In these cases, we expect improvements in precision over the separate analyses similar to the all single camera trap designs, because they are both two sampling methods where identity cannot be deterministically resolved between data sets for any individuals. Given these alternative applications of the SPIM, we suggest the model presented in this paper should be referred to as the 2-flank SPIM.

## APPENDIX A: FULL MCMC ALGORITHM

The joint posterior we want to sample from is

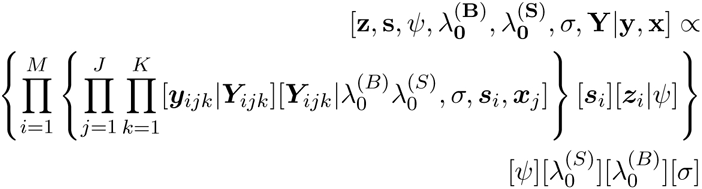

where *M* is the dimension of data augmentation. In practice, the analyst should choose *M >> N* and will need to raise *M* if *N^curr^* = *M* at any point of the MCMC algorithm. The following are our uninformative prior distributions.

1. 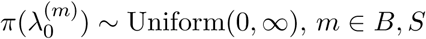
2. *π*(*σ*) ~ Uniform(0, ∞).
3. *π*(*ψ*) ~ Uniform(0, 1)
4. 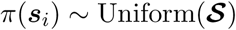

The full conditionals are

1. 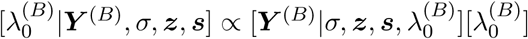, where 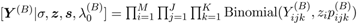
2. 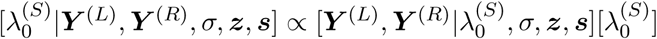, where 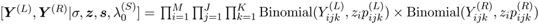
3. 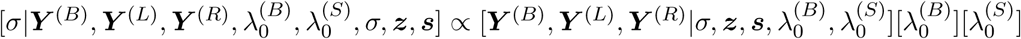, where 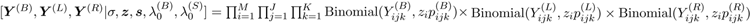
4. 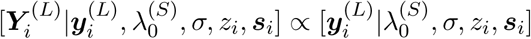, where 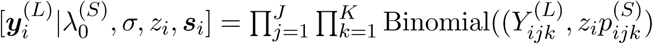
5. 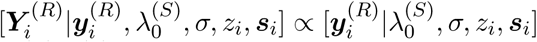, where 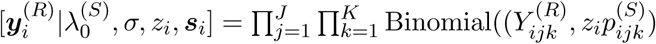
6. 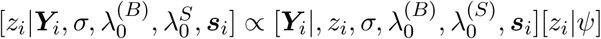, where 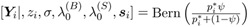 (*p*_*i*_^*^ defined below)
7. 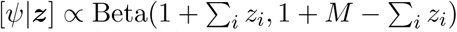
8. 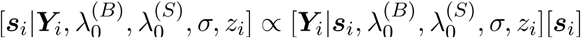, where 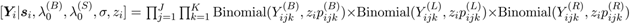

As previously described, conditional on **ID**^(*L*)^ and **ID**^(*R*)^, we can construct a latent true capture history *Y_ijk_* so our MCMC algorithm will follow the standard algorithm as described by Royle et al. (2013) with the additional step of updating **ID**^(*L*)^ and **ID**^(*R*)^ and constructing a new latent true capture history *Y_ijk_* on each MCMC iteration.

1. Update 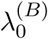 and 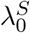 sequentially. Both 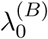 and 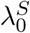 are updated with a Metropolis-Hastings step using the distribution Normal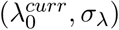 to propose 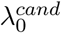, automatically rejecting if a negative value is proposed.
2. Update *σ*. *σ* is updated with a Metropolis-Hastings step using the distribution Normal(*σ^curr^, σ_σ_*), to propose *σ^cand^*, automatically rejecting if a negative value is proposed.
3. Update ***Y*** by updating **ID**^(*L*)^ and **ID**^(*R*)^. On each MCMC iteration, we update both **ID**^(*L*)^ and **ID**^(*R*)^ by swapping *n_swap_* values of **ID**^(*B*)^ stored in **ID**^(*L*)^ and **ID**^(*R*)^. We first update **ID**^(*L*)^. We need to identify the correctly ordered indices **ID**^(*B*)^ at which to swap the value of **ID**^(*L*)^, mapping **ID**^(*L*)^ to **ID**^(*B*)^. We then identify the candidate set of **ID**^(*B*)^ individuals that do not correspond to complete identities (c*_i_* = 0) and who are currently in the population (*z_i_* = 1). From this candidate set, we remove the individuals that would lead to swapping a *z_i_* = 0 individual into the population through the value stored in 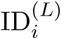. Next, we choose a focal candidate *v* to swap the value of 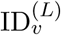 with equal probability across the candidate set. Because proposals that combine candidates whose activity centers are far apart will almost always be rejected, we apply a distance-based criterion to rule out improbable combinations, thus raising acceptance rates. To do this, we calculate the Euclidean distance between the current activity center of the focal candidate *v* and the activity centers of all other individuals in the candidate set. We then identify the set of possible candidate individuals to exchange values of 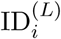 with the focal candidate by identifying which candidate individual activity centers are within a distance threshold, *d_max_*, of the focal individual’s activity center. From this reduced candidate set of size *n_forward_*, we randomly select individual *w* with equal probability 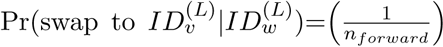 across the remaining candidates and the focal and selected candidate exchange values of 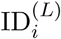. Because this proposal process is not symmetric, we repeat it in reverse to obtain *n_reverse_*, with the probability of choosing this candidate being 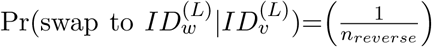. We recompute the proposed true capture history 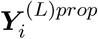 for *i* ∈ *v*, *w* and accept the proposal with probability
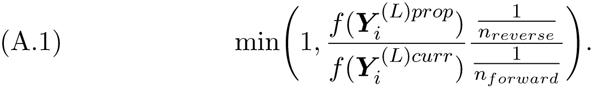

where *f* (.) is the SCR observation model likelihood. This process is then repeated to update **ID**^(*R*)^ and thus, ***Y***^(*R*)^.
4. Update **z**. Each *z_i_* is updated by a Gibbs step using the full conditional above where 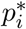 is the probability individual *i* was not captured during the experiment. Let 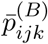 and 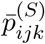 be the probability of not being captured on both and single sides for each individual at each trap on each occasion, respectively. Then 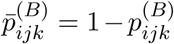 and 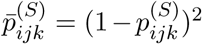 (the squared term is needed because there are two ways to observe a single side capture, right or left side; see model description and trap file sections for definition of 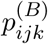 and 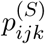 which depend on the number of cameras deployed at each trap and trap operation). The probability of not being captured during the experiment for each individual is then 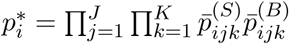.
5. Update *ψ*. *ψ* is updated with a Gibbs step. Since *π*(*ψ*) ~ Uniform(0, 1) is in the Beta family, the full conditional distribution for *ψ* is 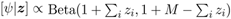
6. Updated **s**. Each activity center ***s**_i_* is updated with a Metropolis-Hastings step using the distributions 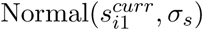 and 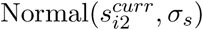 to propose 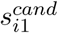 and 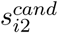, respectively. Proposals that fall outside of the state space are rejected. The full conditional distribution is the SCR observation model likelihood.
7. Record the derived quantities population abundance, 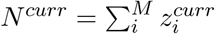, and population density, 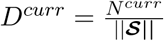

## APPENDIX B: SIMULATIONS

Here, we present a simulation study to assess the performance of the SPIM and compare it to alternative estimators. In addition to the “pragmatic estimators” described in the main article, we will also assess the performance of the “naive independence estimator”. An alternative to the SPIM is to ignore the dependence between the left, right, and both side data sets and average the density estimates from the individual analyses and derive a joint standard error assuming independence. This method is proposed by Wilson, Hammond and Thompson (1999) and while Bonner and Holmberg (2013) point out that assuming independence will lead to the underestimation of standard errors, this estimator might perform reasonably well in some scenarios, such as when data are sparse and thus there is less dependence between data sets. A Bayesian analogue to this method is to perform a joint MCMC analysis on the both (when available), left, and right data sets, allowing each data set to have its own latent structure (***s***,*ψ*,***z***), but sharing detection function parameters (*λ*_0_ and *σ*). On each MCMC iteration, *N_B_* (when both side data is available), *N_L_*, and *N_R_* (current population size values for the both, left, and right side data sets) are independently calculated by summing ***z****_B_*, ***z****_L_*, and ***z****_R_* and their average is recorded.

We conducted 384 simulations for each of 36 scenarios, grouped into four sets, to compare the performance of the SPIM, pragmatic estimators, and the naive independence estimator across a range of trapping array designs and densities. In order to vary the proportion of simulated individuals that produced complete identities, we set 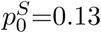, 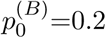 in the first two sets of scenarios and 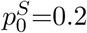, 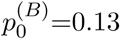 in the second two sets. The first and third sets of scenarios were conducted on a 6 x 6 array and thee second and fourth was conducted on an 8 x 8 array. For all scenarios, *σ*=0.5, trap spacing was 1 unit (2*σ*), and the state space extended 2 units beyond the square trapping arrays in both the X and Y dimensions. The number of IDs to swap on each MCMC iteration, *n_swap_*, was set to 10, and the search radius for activity centers to swap IDs, *d_max_*, was set to 1. Three types of trapping arrays were considered – one with all double camera stations, one with all single camera stations, and a hybrid array with 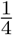 double camera stations and 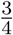 single camera stations (Figure B1). We considered D∈(0.2, 0.4, 0.6) for the 6 x 6 array and density, D∈(0.1, 0.2, 0.4) for the 8 x 8 array. Estimator performance was compared by percent bias of the posterior mode, average mean squared error (MSE), frequentist coverage of the 95% highest posterior density (HPD) intervals, and the mean width of the 95% HPD interval for N. N was chosen over density because the number of individuals to simulate for a given density on the 8 x 8 array of size 121 units^2^ (N=D×121) had to be rounded to the nearest integer so the realized data sets could not be simulated from the exact density. The number of MCMC iterations varied from 35000 to 150000 across scenarios with these numbers chosen to obtain effective sample sizes for *N* greater than 400 and monte carlo standard errors for *N* of less than 0.5.

**Fig B1:**
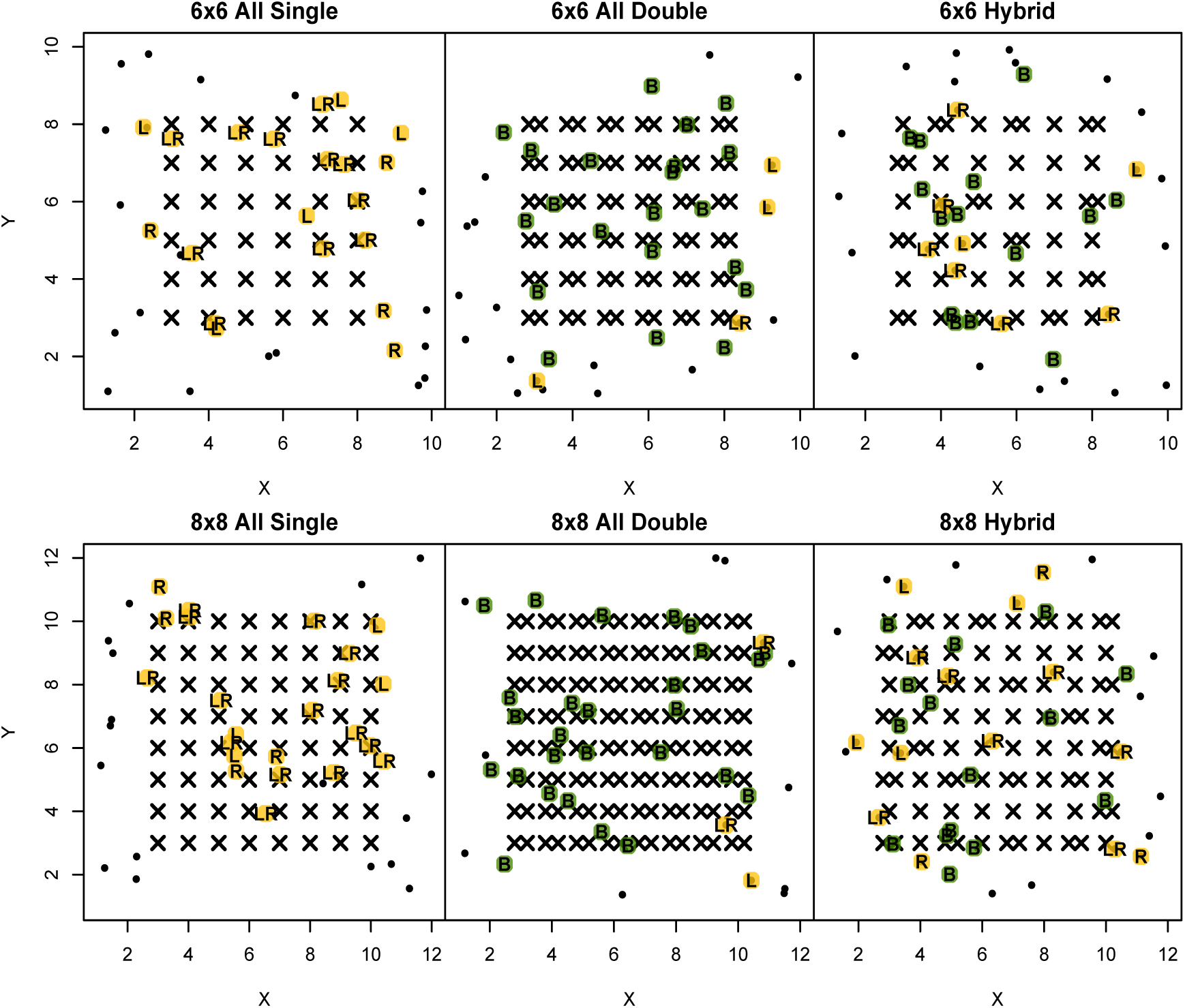
Trapping arrays for the simulation study. Single exes (X) depict single camera stations and double exes (XX) depict double camera stations. Activity centers from one realization of the capture process are displayed, with green dots representing complete identity individuals (B), yellow dots representing partial identity individuals captured on the left side (L), right side (R) or left and right side (LR). Black dots representing individuals never captured.

In the scenarios where data are more sparse, occasionally there were realizations of the capture process that did not produce a spatial recapture – a capture of the same animal at more than one location. Analyzing data sets with no spatial recaptures leads to density estimates that are biased high (Sun, Fuller and Royle, 2014); therefore, for simulated data sets with no spatial recaptures, data sets were discarded. For simulated data sets with spatial recaptures between the three data sets, but not within the single side or both plus single side data sets, the single side estimators were not fit. For simulated data sets that did not have spatial recaptures in all two or three data sets, the naive independence estimator was not fit. In our simulations, the only way to obtain a complete identity was by being captured on both sides simultaneously at least once during the survey. We used linear regression on the response variable of mean difference in 95% credible interval widths between the SPIM and best-side estimators to test the hypotheses that precision gains in the SPIM are related to the mean number of complete identity individuals captured and the percentage of captured individuals with complete identities. Finally, the R package stargazer was used to convert all R output to LaTeX tables (Hlavac, 2015).

### B.1. Simulation Results

For all single camera trapping arrays, the random-side estimator produced nearly unbiased density estimates (Figure B2), while the best-side estimator was biased high roughly 5% when 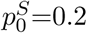 and roughly 15% when 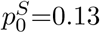. The SPIM was biased high, but less than 5%, except for the scenario with the lowest population size where it was biased low by 7%. Coverage for these three estimators was roughly nominal or above nominal. On average, the SPIM decreased the 95% HPD interval width by 30-40% with larger increases at smaller population sizes and when 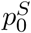 was lower (Figure A3a). The SPIM decreased the MSE by 40-60% over the best-side estimator (Figure A3a) and the random-side estimator (see Supplement A). The naive independence estimator was generally biased high (up to 12.8%), and bias decreased as *N* increased (see Supplement A for naive independence estimator results). Coverage for the naive independence estimator was slightly less than nominal and the mean width of the 95% HPD interval was larger than that of the SPIM except in some of the scenarios where 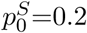 and *N* was larger; however coverage in these scenarios was around 0.90.

**Fig B2:**
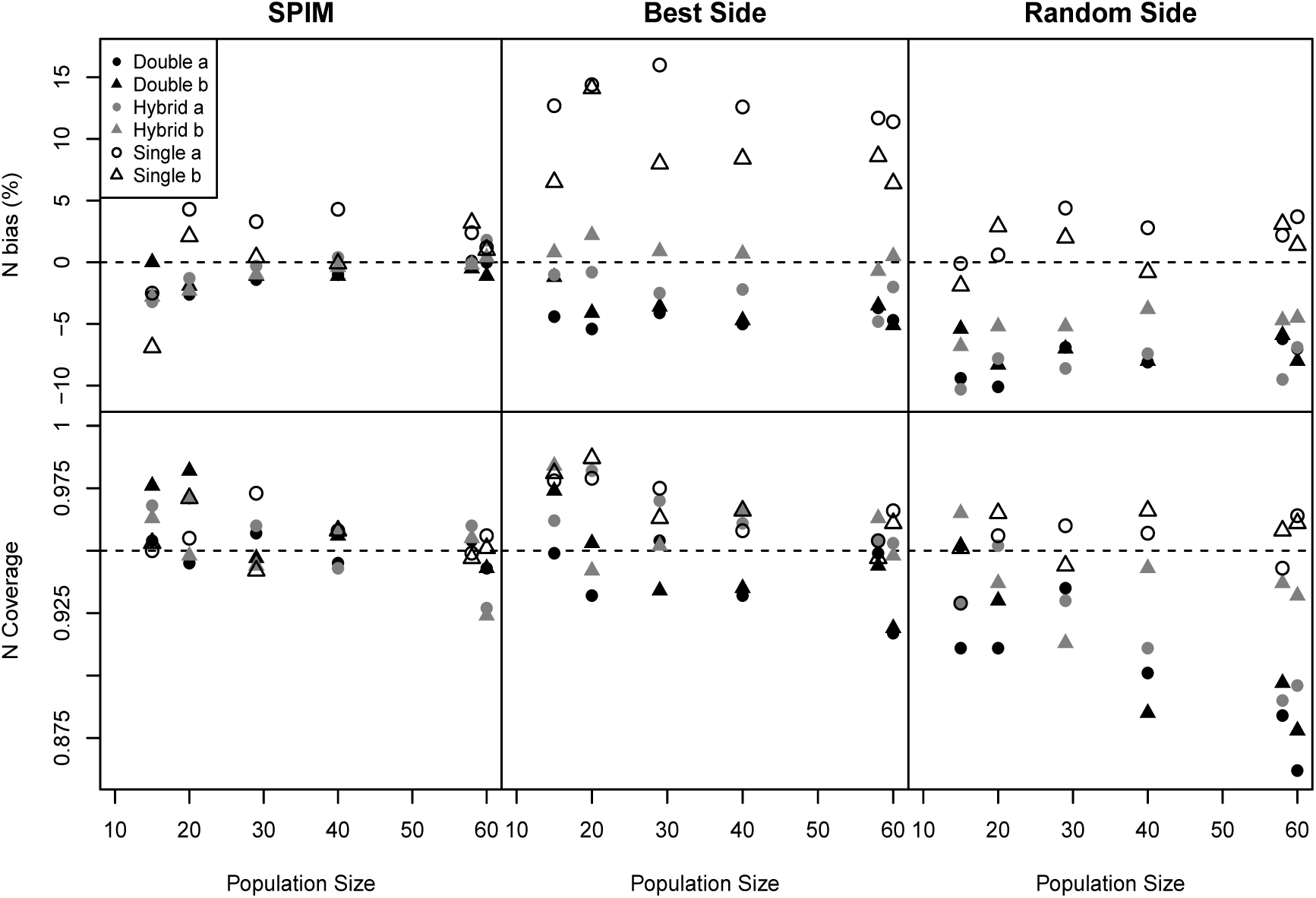
Bias and coverage of population size for the SPIM, best-side, and random-side estimators. Scenarios labeled “a” correspond to scenarios with 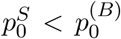 and those labeled “b” correspond to scenarios with 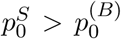. Double indicates two camera per station, single indicates one camera per station, and hybrid indicates a combination of double and single stations as depicted in Figure A1.

For all double camera trapping arrays, the both-plus-random-side and both-plus-best-side estimators were biased low 5-7% (Figure B2) due to the individual heterogeneity induced when constructing these data sets, but the both-plus-best-side was less biased because always choosing the best side induces positive bias as seen in the single camera simulations, counteracting the negative bias from ignored individual heterogeneity. The SPIM had a slight negative bias that disappeared as *N* increased. The both-plus-best-side estimator had nominal coverage at low *N*, but coverage tended to be less than nominal as *N* increased. The both plus random-side estimator had lower than nominal coverage that decreased with *N*. The SPIM had nominal or greater than nominal coverage. On average, the SPIM produced 95% HPD intervals that were of equal size or slightly wider (4%) than the best-side estimator. (Figure B3a). The SPIM produced point estimates with slightly lower MSE, with a greater improvement at larger *N*. The naive independence estimator was biased high, but less so than in the all single trapping array scenarios, and bias decreased with increasing *N*. Coverage for the naive independence estimator was around 0.85 in all scenarios and the mean width of the 95% HPD interval was similar to that of the SPIM and single-side estimators.

**Fig B3:**
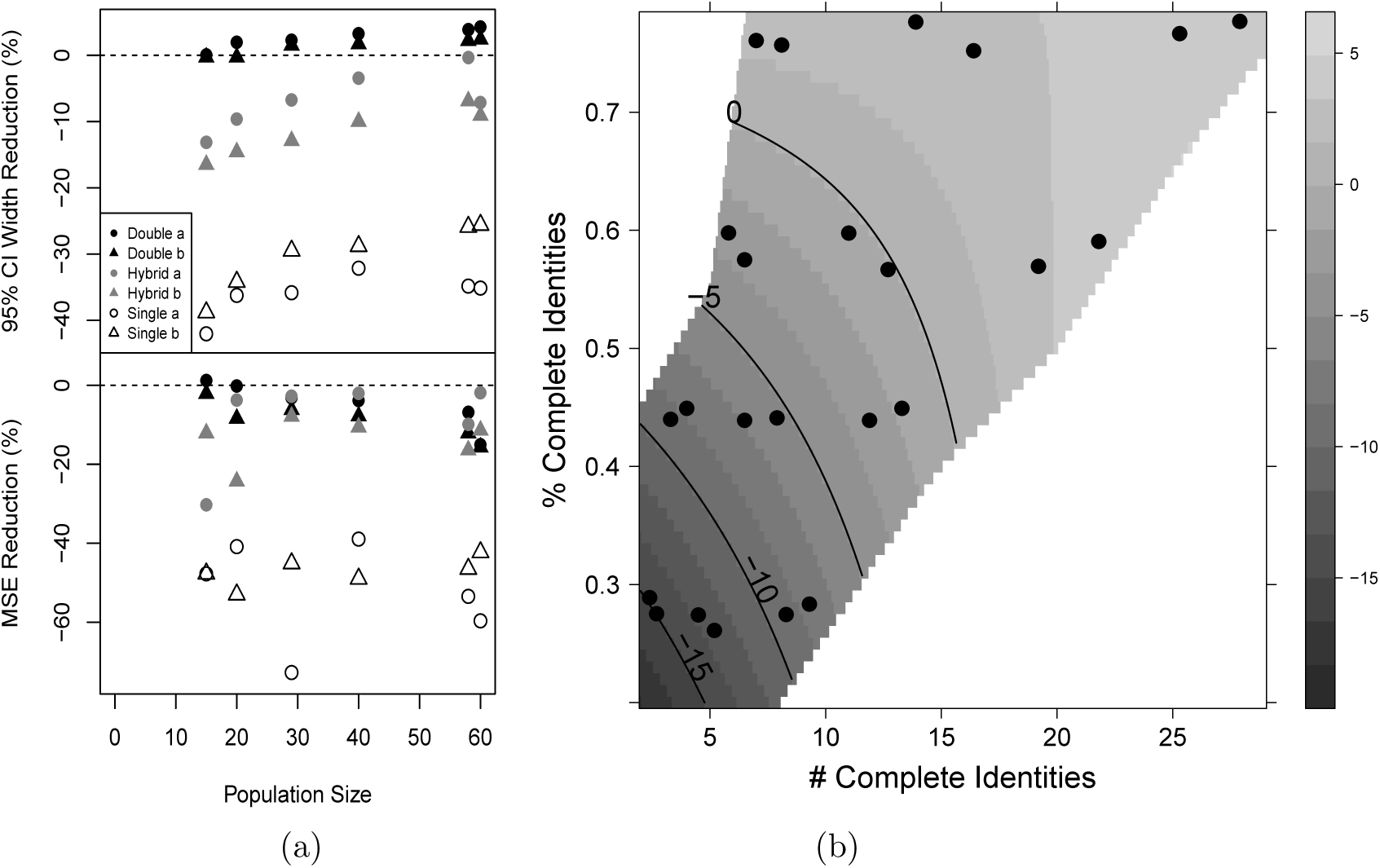
(a) Performance difference between the SPIM and best-side estimator as judged by the mean reduction in the width of the 95% credible interval and the mean reduction in MSE. Scenarios labeled “a” correspond to scenarios with 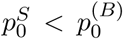 and those labeled “b” correspond to scenarios with 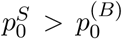. (b) The mean difference in the 95% credible interval width between the SPIM and best side estimator by the mean number of complete identity individuals captured and the mean percentage of captured individuals that had complete identities. The scenarios with *>*50% complete identities are the all double camera scenarios and those with *<*50% complete identities are the hybrid scenarios and the % complete identities are higher when 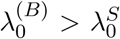. Within each scenario, the number of complete identities increase as *N* increases.

For hybrid camera trapping arrays, the both-plus-single-side estimators exhibited the same patterns as in the all double camera trapping arrays, but to a lesser degree. The both-plus-random-side estimator was still biased low, but the both-plus-best-side estimator was now unbiased due to the two sources of bias roughly canceling out (Figure B2). Coverage for the both-plus-best-side estimator was nominal or higher and coverage for the both-plus-random-side was less than nominal except at the lowest *N*. The SPIM performed about the same in terms of bias and coverage as it did in the all double trap scenarios. On average, the SPIM produced 95% HPD intervals that were 5-17% more narrow than the both-plus-best-side estimator. (Figure B3a), with the largest precision gains seen when *N* was lower. MSE reductions were similar to the all double trap scenarios. The difference in precision between the SPIM and best-side estimator was related to the mean number of complete identity individuals captured, the percent of captured individuals whose identity was complete, and their interaction (all p*<*0.0001). The number of complete identity individuals influenced precision more when the percent of individuals whose identity was complete was lower (Figure B3b). The naive independence estimator was biased high when 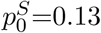 as much as 20% but moderately biased low when 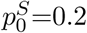. Coverage for the naïve independence estimator was slightly less than nominal in all scenarios and the mean width of the 95% HPD interval was larger than that of the SPIM and single-side estimators.

In the lowest density simulations on all single camera trapping arrays when 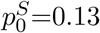, 14-20% of the simulated data sets did not have spatial recaptures within either the best-side or random-side data sets and therefore were excluded from these analysis. In practice, one could deviate from the best-side or random-side rule if the other data set had a spatial recapture, but the SPIM was able to accommodate the realizations with spatial recaptures between, but not within data sets while maintaining acceptable bias and nominal coverage. Full simulation results can be found in Supplement A.

### B.2. Simulation Discussion

When using all single camera trap stations, the best-side estimator was significantly biased high and although the random-side-estimator is unbiased, the SPIM was significantly more precise and accurate (see Supplement A). The difference in precision between the SPIM and the random-side estimator was similar to the best-side comparisons in Figure A3a and MSE reductions were moderately less than the best-side comparisons due to the lack of bias in the random-side analysis. When at least some double camera trap stations are used and thus some identities are complete, aggregating the single-side capture histories for the complete identity individuals introduced individual heterogeneity in capture probability and thus negative bias and reduced coverage into the single-side analyses. For the best-side estimator, the positive bias due to always selecting the data set with the most individuals was roughly canceled out by the negative bias from individual heterogeneity in the hybrid trapping array designs; however, it is not likely this will hold across all combinations of parameter values. The best-side estimator was biased low in the double camera trapping array designs, suggesting that performance depends on the ratio of complete to partial identity individuals, which determines the magnitude of individual heterogeneity in capture probability. The SPIM had minimal bias and nominal coverage in the hybrid and double trapping array designs and we expect this to hold across a wide range of parameter values and trapping array designs. Precision of the SPIM was slightly less than the best-side estimator in some of the double camera trapping array designs; however, coverage of the best-side estimator in most of these scenarios was slightly less than nominal. In the hybrid designs with fewer complete identity individuals, the SPIM moderately increased precision and reduced MSE. The performance gain of the SPIM is further increased when considering other options available to the researcher. If both data sets were analyzed rather than just the best or random-side, the researcher could choose either the most precise estimate, a protocol that will guarantee less than nominal coverage, or the most conservative estimate in which case the precision gains of using the SPIM will be increased. The naive independence estimator was biased high in all scenarios except when 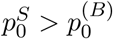 on hybrid trapping arrays, exhibited slightly to moderately low coverage, and was not more precise than the SPIM except in a few scenarios with the most captured individuals. If the goal is to maintain good frequentist properties, researchers should choose the analysis method before examining their data and we argue that the SPIM is the best all-around choice to achieve these ends.

### SUPPLEMENTARY MATERIAL

**Supplement A: Simulation Tables for Appendix A** (Submitted to AOAS).

**Supplement B: Comparison of Spatial Partial Identity Model to the Non-spatial Partial Identity Model** (Submitted to AOAS).

